# Leveraging prior biological knowledge improves prediction of tocochromanols in maize grain

**DOI:** 10.1101/2022.08.16.502005

**Authors:** Ryokei Tanaka, Di Wu, Xiaowei Li, Laura E. Tibbs-Cortes, Joshua C. Wood, Maria Magallanes-Lundback, Nolan Bornowski, John P. Hamilton, Brieanne Vaillancourt, Xianran Li, Nicholas T. Deason, Gregory R. Schoenbaum, C. Robin Buell, Dean DellaPenna, Jianming Yu, Michael A. Gore

## Abstract

With an essential role in human health, tocochromanols are mostly obtained by consuming seed oils; however, the vitamin E content of the most abundant tocochromanols in maize grain is low. Several large-effect genes with *cis*-acting variants affecting mRNA expression are mostly responsible for tocochromanol variation in maize grain, with other relevant associated quantitative trait loci (QTL) yet to be fully resolved. Leveraging existing genomic and transcriptomic information for maize inbreds could improve prediction when selecting for higher vitamin E content. Here, we first evaluated a multikernel genomic best linear unbiased prediction (MK-GBLUP) approach for modeling known QTL in the prediction of nine tocochromanol grain phenotypes (12–21 QTL per trait) within and between two panels of 1,462 and 242 maize inbred lines. On average, MK-GBLUP models improved predictive abilities by 7.0 to 13.6% when compared to GBLUP. In a second approach with a subset of 545 lines from the larger panel, the highest average improvement in predictive ability relative to GBLUP was achieved with a multi-trait GBLUP model (15.4%) that had a tocochromanol phenotype and transcript abundances in developing grain for a few large-effect candidate causal genes (1–3 genes per trait) as multiple response variables. Taken together, our study illustrates the enhancement of prediction models when informed by existing biological knowledge pertaining to QTL and candidate causal genes.

**Core Ideas:**

- With varying levels of vitamin E activity, tocochromanols found in maize grain are essential for human health
- Selecting for higher vitamin E content in maize grain can be enhanced with genomic prediction
- Prediction models leveraging existing biological knowledge were evaluated in two panels of maize inbred lines
- Multikernel prediction models based on previously identified QTL improved predictive ability
- A multi-trait prediction model that had transcript abundances of a few large-effect causal genes performed the best

## INTRODUCTION

Tocopherols and tocotrienols, together termed tocochromanols, are plant-synthesized lipid-soluble antioxidants that are most abundant in seed where they prolong seed viability during desiccation and storage by limiting lipid peroxidation (Sattler et al., 2004; Mène-Saffrané et al., 2010). The rings of the four different types of tocopherols and tocotrienols (α, β, δ, and γ) differ in their numbers and positions of methyl groups. In mammals, tocochromanols are an essential dietary nutrient as vitamin E and while most tocochromanols have some level of vitamin E activity, α-tocopherol (αT) has the highest activity and is preferentially retained from the diet in humans (DellaPenna & Mène-Saffrané, 2011). Though tocotrienols have lower vitamin E activity than their corresponding tocopherols, their more even distribution in membranes makes them better at scavenging lipid peroxyl radicals than αT (Jiang, 2014). The low level of tocochromanols with high vitamin E activity, especially αT, in the seed oils of most cereal crops contributes to the suboptimal daily intake of vitamin E in human diets (DellaPenna & Mène-Saffrané, 2011).

Maize is one of the most important staple food crops worldwide. Unfortunately, the predominant maize varieties grown around the world are inadequate in providing grain that meets the recommended dietary intake of vitamin E for healthy individuals (Fitzpatrick et al., 2012). Considerable natural variation, however, exists for levels of tocopherols and tocotrienols in maize grain (Li et al., 2012; Lipka et al., 2013; Baseggio et al., 2019; Wu et al., 2022), thus there is tremendous potential for maize biofortification breeding efforts centered on the elevation of vitamin E and antioxidants levels in maize grain (Diepenbrock & Gore, 2015; Mène-Saffrané & Pellaud, 2017). To identify causal genes controlling grain tocochromanol levels, most genome-wide association studies (GWAS) used moderately-sized maize mapping panels with low genotyping densities that limited identification to a few large-effect loci involved in the genetic control of these nutritionally important grain phenotypes (Li et al., 2012; Lipka et al., 2013; Wang et al., 2018; Baseggio et al., 2019). The majority of these large-effect loci shared across these studies encode core tocochromanol pathway enzymes (γ-tocopherol methyltransferase, *vte4*; tocopherol cyclase, *vte1*; and homogentisate geranylgeranyltransferase, *hggt1*), but they do not fully explain the phenotypic variance for grain tocochromanol phenotypes (Li et al., 2012; Lipka et al., 2013; Wang et al., 2018; Baseggio et al., 2019).

To more fully elucidate the genetic architecture of tocochromanol levels in maize grain, a joint-linkage analysis and GWAS was conducted in the 5,000-line U.S. maize nested association mapping (NAM) panel scored with ∼29 million sequence variants (Diepenbrock et al., 2017). Diepenbrock et al. (2017) identified 154 joint-linkage quantitative trait loci (JL-QTL) for individual and total tocochromanols (12–21 JL-QTL/phenotype) that reduced to a nonredundant set of 50 QTL, of which 13 were resolved to mostly large-effect single genes. Of the identified 13 candidate causal genes, seven encode enzymes involved in the synthesis of tocochromanols or their precursors (1-deoxy-D-xylulose-5-phosphate synthase, *dxs2*; solanesyl-DP synthase, *sds*; arogenate dehydrogenase, *arodeH2* Zm00001d014734; p-hydroxyphenylpyruvate dioxygenase, *hppd1*; 2-methyl-6-phytyl-1,4-benzoquinol/2-methyl-6-geranylgeranyl-1,4-benzoquinol methyltransferase, *vte3*; *hggt1*; and *vte4*), whereas six code for activities novel to tocochromanols (protochlorophyllide reductase, *por1* and *por2*; snare protein, *snare*; lipid transfer protein, *ltp*; PHD finger transcription factor, *phd*; and fibrillin, *fbn*). Implicating the importance of gene expression variation, seven of the 13 genes (*dxs2*, *hggt1*, *vte4*, *por1*, *por2*, *phd*, and *fbn*) were determined to be correlated expression and effect QTL (ceeQTL, Diepenbrock et al., 2021), because the expression level of each gene was significantly associated with the allelic effect estimates of its respective trait-specific JL-QTL at multiple time points of kernel development.

In a transcriptome-wide association study (TWAS), mRNA expression levels are correlated with terminal phenotypes, providing a high-resolution, gene-based association test that is less influenced by LD patterns, as opposed to DNA genetic markers in GWAS (Hirsch et al., 2014; Pasaniuc & Price, 2016; Lin et al., 2017; Kremling et al., 2019; Li et al., 2021; Hershberger et al., 2022). Through combining GWAS and TWAS with the Fisher’s combined test in the maize Ames panel of nearly 1,500 inbred lines, Wu et al. (2022) further elucidated the genetic architecture of grain tocochromanol phenotypes that had not been fully resolved in the NAM panel, identifying novel associations with an *arodeH2* paralog (Zm00001d014737), *dxs1*, *vte5* (phytol kinase), *vte7* (α-/β-hydrolase family protein), and *samt1* (S-adenosylmethionine transporter), along with the re-identification of *por1*, *por2*, *dxs2*, *arodeH2* Zm00001d014734, *hppd1*, *vte1*, *vte4*, and *hggt1*. Of these 13 genes, seven were detected by TWAS (*por1*, *por2*, *dxs1*, *dxs2*, *vte4*, *hggt1*, and *samt1*), thus further demonstrating that regulatory variation plays an important role in influencing tocochromanol levels in maize grain. However, a modest number of small- to moderate-effect NAM JL-QTL remained unresolved, but are still relevant to genomics-assisted breeding approaches.

Genomic selection uses a prediction model trained on genome-wide markers and phenotypes to predict genomic estimated breeding values for target set individuals that have only marker genotype data (Lorenz et al., 2011; Heslot et al., 2015). The genomic best linear unbiased prediction (GBLUP) model is one of the most commonly used models for genomic prediction (GP) of complex trait variation with genome-wide markers, given that it assumes polygenic inheritance—many loci with small effect sizes (Meuwissen et al., 2001). However, for phenotypes having less complex genetic architectures that include large-effect QTL like tocochromanol levels in maize grain, Bayesian regression models (*e.g.*, BayesA and BayesB) that assume the existence of loci with major effects could achieve higher predictive performance than GBLUP (de Los Campos et al., 2013; Gianola, 2013). Illustrative of the potential for genomic selection, a range of low to moderately high predictive abilities were obtained via GP of tocochromanol levels in fresh kernels of a sweet corn diversity panel (GBLUP model; Baseggio et al., 2019; Hershberger et al., 2022) and in mature maize grain samples of both the Ames panel and an exotic derived population (GBLUP and Bayesian models; Tibbs-Cortes et al., 2022)

In addition to Bayesian models, it is possible to improve model performance for phenotypes predominantly controlled by a few large-effect loci through introducing significant trait-associated markers as fixed covariates in the GBLUP model (e.g., Bernardo, 2014; Spindel et al., 2016). Relatedly, there have been attempts to enhance trait prediction models by including a genomic relationship matrix (GRM) constructed with a subset of genetic markers surrounding strong association signals (Zhang et al., 2014; Li et al., 2018; Liu et al., 2019), or *a priori* candidate genes from trait-relevant biosynthetic pathways (Owens et al., 2014; Baseggio et al., 2019, 2020). The latter of which, however, resulted in lower predictive abilities relative to genome-wide markers for tocochromanol levels in fresh sweet corn kernels (Baseggio et al., 2019). In contrast to a single GRM approach, multikernel models (Speed & Balding, 2014) are a promising yet unexplored prediction approach to account for the large-effect loci associated with tocochromanol grain phenotypes, as separating genetic markers into functional (causal or potentially causal genes) and nonfunctional sets through the calculation of multiple GRMs has improved predictions for plant metabolic phenotypes (Turner-Hissong et al., 2020; Campbell et al., 2021a; b; Brzozowski et al., 2022).

The inclusion of molecular intermediates such as mRNA expression levels in prediction models could strengthen connections between genotype and terminal phenotypes (e.g., Riedelsheimer et al., 2012; Guo et al., 2016; Schrag et al., 2018; Azodi et al., 2020; Hu et al., 2021). Although dependent on the target phenotype, transcriptome-based prediction can achieve predictive abilities that are comparable to those of prediction with solely genetic markers (Guo et al., 2016; Schrag et al., 2018; Azodi et al., 2020). When implemented for the prediction of tocochromanols and carotenoids in fresh sweet corn kernels, Hershberger et al. (2022) showed models that combined transcriptome data from developing kernels with genome-wide markers generally had higher predictive abilities relative to models with only markers or transcript abundances. Taken together, these results support the need for further research endeavors that leverage transcriptome data to harness regulatory variation for predicting tocochromanols in mature maize grain.

Tocochromanol maize grain phenotypes are model phenotypes for the evaluation of genome- and transcriptome-based prediction approaches that leverage prior biological information. The objectives of this study as it relates to the prediction of tocochromals in maize grain were to (i) assess GP models with multiple kernels that separate single-nucleotide polymorphism (SNP) genetic markers according to JL-QTL identified in the maize NAM panel, (ii) evaluate transcriptome-based prediction models varying in the degree to which they prioritize candidate causal genes, and (iii) reveal the candidate causal genes that are of highest importance to model performance.

## MATERIALS AND METHODS

### Phenotype data processing

The first data set of tocochromanol phenotypes was generated by Wu et al. (2022). Briefly, physiologically mature self-pollinated ears were harvested from 1,815 inbred lines of the maize North Central Regional Plant Introduction Station panel (hereinafter the Ames panel, Romay et al., 2013) that was grown in an augmented complete block design during the 2015 and 2017 field seasons at Iowa State University Agricultural Engineering and Agronomy Research Farm in Boone, IA. For each harvested plot, bulked grain samples from shelled ears were ground and analyzed for tocopherol and tocotrienol concentrations via high-performance liquid chromatography (HPLC) and fluorometry as previously described (Lipka et al., 2013). Through a mixed linear model analysis, Wu et al. (2022) generated outlier-screened best linear unbiased estimator (BLUE) values in μg g^−1^ dry seed for nine phenotypes (α-tocopherol, αT; δ-tocopherol, δT; γ-tocopherol, γT; α-tocotrienol, αT3; δ-tocotrienol, δT3; γ-tocotrienol, γT3; total tocopherols, ΣT: αT + δT + γT; total tocotrienols, ΣT3: αT3 + δT3 + γT3; and total tocochromanols, ΣTT3: total tocopherols + total tocotrienols) scored on 1,462 inbred lines that were not classified as sweet corn, popcorn, or having an endosperm mutation. Next, Wu et al. (2022) transformed the BLUE values of each phenotype with the Box-Cox procedure. These transformed BLUE values served as the Ames panel phenotypic data set.

The second data set of tocochromanol phenotypes was initially generated by Lipka et al. (2013) through identical HPLC-based quantification of tocopherols and tocotrienols in ground mature grain samples prepared from the maize Goodman panel (also known as the maize 282 association panel, Flint-Garcia et al., 2005; Gage et al., 2020) that had been grown in an incomplete block α-lattice design at Purdue University Agronomy Center for Research & Education in West Lafayette, IN, in 2009 and 2010. The mixed linear model analysis implemented by Lipka et al. (2013) produced outlier-screened best linear unbiased predictor (BLUP) values in μg g^−1^ dry seed for the same nine grain tocochromanol phenotypes scored on the 252 inbred lines. To be consistent with the Ames panel phenotypic data set, we removed lines classified as sweet corn, popcorn or having an endosperm mutation, resulting in the retention of 242 lines with Box-Cox transformed BLUP values that comprised the phenotypic data set of the Goodman panel.

### Genotype data processing and imputation

We constructed a high-density SNP marker set in B73 RefGen_v4 coordinates for the Ames and Goodman panels from target and reference SNP genotype sets following the marker genotype imputation approach of Wu et al. (2021). The downloaded (ZeaGBSv27_publicSamples_raw_AGPv4-181023.vcf.gz, available at http://datacommons.cyverse.org/browse/iplant/home/shared/panzea/genotypes/GBS/v27) data set of 943,455 unimputed genotyping-by-sequencing (GBS) SNPs called by Romay et al. (2013) served as the foundation of the target SNP genotype set. Each panel had its own sequencing project, thus GBS samples for each inbred line are specific to either the Ames or Goodman panel. To generate a target SNP set for the Goodman panel, unique GBS samples for each of the 242 lines were identified from GBS samples annotated as the “Ames282” project. Next, we filtered for biallelic SNPs that were polymorphic among the 242 lines, resulting in a set of 361,110 GBS SNPs. The target SNP set of 443,419 GBS SNPs for the 1,462 lines of the Ames panel was generated by Wu et al. (2022) in an identical approach that included a consensus genotype calling step to merge two or more GBS samples from the same line. Our implemented consensus genotype calling step was modified from an approach developed by Dzievit et al. (2021). By taking the intersection of SNPs between panels, we generated a shared target set of 349,871 unimputed biallelic GBS SNPs that were scored across all 1,704 lines of the Ames and Goodman panels. Prior to imputation, we set all heterozygous genotype calls to missing. The reference marker genotype set of 14,613,169 SNPs constructed from maize HapMap 3.2.1 (Bukowski et al., 2018) by Wu et al. (2021) was imputed with BEAGLE v5.0 (Browning et al., 2018) based on the GBS SNPs (target set) in the combined set of 1,704 lines. All parameters for imputation with BEAGLE v5.0 were identical to those used in Wu et al. (2021). To generate a set of high quality imputed genotypes, we subsetted 12,045,820 biallelic SNPs with minor allele frequency (MAF) ≥ 1% and predicted dosage *r^2^* (DR2) ≥ 0.80. This set of 12,045,820 SNPs was linkage disequilibrium (LD) pruned (pairwise *r*^2^ < 0.10) in PLINK version 1.9 (Purcell et al., 2007) with a sliding window of 100 SNPs and step size of 25 SNPs, resulting in a reduced marker set of 341,189 SNPs for 1,704 lines that was used for our implemented computationally intensive statistical analyses.

### Expression data processing

The developing kernel gene expression data set was produced as previously described (Wu et al., 2022). In brief, 1,023 of the 1,815 inbred lines from the Ames panel and five additional parents of the U.S maize NAM panel (Yu et al., 2008; McMullen et al., 2009) were evaluated in an augmented incomplete block design during the 2018 field season at Iowa State University Agricultural Engineering and Agronomy Research Farm in Boone, IA. From each plot, a single self-pollinated ear was hand-harvested at ∼23 days after pollination, with frozen kernels from the mid-section of the ear ground for RNA isolation using a modified hot borate protocol (Wan & Wilkins, 1994). Isolated RNA samples were used to construct libraries with the Lexogen QuantSeq 3′ mRNA-Seq Library Kit FWD (Lexogen, Greenland, NH). Libraries were then sequenced on an Illumina NextSeq 500 to 85 nt in single end mode (Illumina, San Diego, CA) at the Genomics Facility of the Cornell Institute of Biotechnology. The 3′ QuantSeq reads were cleaned with Cutadapt version 2.3 (Martin, 2011), aligned to the B73 RefGen_v4 reference genome (Jiao et al., 2017) using HISAT2 version 2.1.0 (Kim et al., 2019), and alignments sorted with SAMTools version 1.9 (Li et al., 2009). Next, read counts were generated using the htseq-count function within HTSeq version 0.11.2 (Anders et al., 2015) based on the B73 version 4.59 annotation and then normalized using the *rlog* function in the DESeq2 package (Love et al., 2014). After the implementation of several additional stringent quality control measures at the sample and gene levels as described in Wu et al. (2022), a total of 741 high quality samples from check and noncheck lines with *rlog*-transformed read counts for 22,136 genes were retained for generating BLUE values of each gene by fitting a mixed linear model. To coincide with both tocochromanol phenotype data sets, 545 lines were retained after removing 104 lines classified as sweet corn, popcorn, or an endosperm mutant. Lastly, we included the expression BLUE values of the *vte7* locus that were generated with a slightly modified pipeline to account for tandemly duplicated genes (Zm00001d006778 and Zm00001d006779) in B73RefGen_v4 as described in Wu et al. (2022). In total, the generated data set consisted of expression BLUE values for 22,137 gene loci profiled on 545 lines.

### Genomic prediction

We evaluated two GP models with genome-wide SNP markers, GBLUP and BayesB (Meuwissen et al., 2001; Gianola, 2013; Pérez & de los Campos, 2014). BayesB uses a unique prior distribution that strongly shrinks the estimated marker effects toward zero (Gianola, 2013), thus it is an ideal model to be compared with GBLUP (Heslot et al., 2012). The LD-pruned data set of 341,189 SNPs scored on the 1,704 lines was used for both GP models as a compromise between computational efficiency and model performance. The GRM for GBLUP was calculated based on VanRaden’s method 1 (VanRaden, 2008) from the 341,189 SNPs. With each Box-Cox transformed phenotype as a response variable, the two GP modeling approaches were conducted to assess mean predictive ability of 10 iterations from 5-fold cross-validation (CV) within the Ames and Goodman panels in the BGLR package version 1.0.8 (Pérez & de los Campos, 2014) in R version 3.5.0 (R Core Team, 2021) with a Markov chain Monte Carlo (MCMC) process having 12,000 iterations with a burn-in of 8,000 and a thinning of 5. Predictive ability was estimated as the Pearson’s correlation (*r*) of back-transformed genomic estimated breeding values and untransformed BLUE or BLUP values.

We also assessed predictive ability with the Ames panel as the training population and the Goodman panel as the prediction population and vice versa in the BGLR package version 1.0.8 with an MCMC process that had 60,000 iterations with a burn-in of 40,000 and a thinning of 20. To conduct predictions between the Goodman and Ames panels, we first had to exclude lines overlapping between both panels, so as not to overestimate predictive abilities. Briefly, the 341,189 SNPs were used to calculate pairwise identity-by-state (IBS) between lines, with pairs of lines considered to be overlapping between panels if they had an IBS value ≥ 0.8 and shared the identical or nearly identical inbred line name. Of the 242 lines in the Goodman panel, 169 lines were found to overlap with the Ames panel (Supplemental Table S1). Within the Ames panel, 10 of the 169 lines were duplicated according to their inbred line name but each had a different accession number. These 10 ‘line name duplicates’ had an IBS value ≥ 0.8 with a line of the same name in the Goodman panel, thus they too were labeled as overlapping lines. Separately, we found in the Ames panel that one of two lines with the same name but a different accession number had an IBS value ≥ 0.8 with a same-named line in the Goodman panel, thus both this one high IBS line in the Ames panel and the corresponding line of the same name in the Goodman panel were designated to be overlapping. Therefore, in these two prediction scenarios, the Goodman and Ames panels consisted of 73 and 1,283 non-overlapping lines, respectively, when serving as the prediction population.

To incorporate the 12–21 NAM JL-QTL associated with each of the nine tocochromanol phenotypes (Diepenbrock et al., 2017) that had been uplifted to B73 RefGen_v4 coordinates by Wu et al. (2022) into GP models, we used a multikernel GBLUP (MK-GBLUP) model in the BGLR package version 1.0.8 (Pérez & de los Campos, 2014) that partitioned genetic effects into multiple random effects (Supplemental Table S2). The MCMC parameters used for MK-GBLUP in the four GP scenarios (within Ames, within Goodman, from Ames to Goodman, and from Goodman to Ames) are the same as those employed for GBLUP and BayesB. For each Box-Cox transformed tocochromanol phenotype, a covariance matrix for each JL-QTL (hereinafter, we refer to this covariance matrix as a QTL-specific relationship matrix) was calculated based on VanRaden’s method 1 (VanRaden, 2008) with the following marker sets subsetted from the 341,189 SNPs: 1) SNPs ± 250 kb from the JL peak SNP marker, (2) SNPs ± 1 Mb from the JL peak SNP marker, or (3) SNPs within the JL-QTL support interval (SI). After defining the QTL-specific relationship matrices for each phenotype, a GRM was calculated with the remaining unincluded SNPs. All covariance matrices for each phenotype were additively included in the MK-GBLUP model. In addition to the evaluation of the predictive ability of the MK-GBLUP model for each phenotype in the four GP scenarios as performed for GBLUP and BayesB, the predictive ability of each kernel was assessed by partial correlation analysis with the ppcor package in R version 3.5.0 (R Core Team, 2021).

### Transcriptome-based prediction

We conducted transcriptome-based prediction of the nine Box-Cox transformed tocochromanol phenotypes in the 545 lines of the Ames panel with expression data at the transcriptome-wide and pathway-level. The construction of transcriptomic relationship matrices (TRMs) from expression-BLUE values of genes was performed based on a linear kernel as previously described (Hershberger et al., 2022). The transcriptome-wide TRM (TRM.all) was constructed from the expression-BLUE values of all 22,137 gene loci, whereas the pathway-level TRM (TRM.cand) was constructed from the subsetted expression-BLUE values of 111 of a possible 126 *a priori* candidate gene loci encoding activities pertaining to the biosynthesis of tocochromanols in maize grain (Supplemental Table S3). The GRM for this prediction analysis was constructed based on VanRaden’s method 1 (VanRaden, 2008) from 341,034 of the 341,189 SNPs that were polymorphic among the 545 lines. We evaluated the mean predictive ability of 10 iterations from 5-fold CV of five models that had different combinations of relationship matrices (Table 1), as well as the GBLUP model that only uses a GRM of the 545 lines in the BGLR package version 1.0.8 (Pérez & de los Campos, 2014) with MCMC parameters identical to those used for the GP with CV. For the two GRM+TRM models, regression coefficients (*i.e.*, estimated effects) of transcripts were calculated based on the method described in Zhang et al. (2021). To perform this calculation, we fit the model on all 545 lines with an MCMC process that had 60,000 iterations with a burn-in of 40,000 and a thinning of 20. The estimated regression coefficients for the six tocochromanol compounds (αT, δT, γT, αT3, δT3, and γT3) were analyzed with a principal component analysis (PCA) using *prcomp* function from the R base package (R Core Team, 2021) with standardization to visualize genes with large estimated effects on the phenotypic variation of tocochromanols.

**Table 1.** Summary of the evaluated transcriptome-based prediction models

| Model <sup>a</sup> | No. gene loci with transcript abundances | Response variable | Relationship matrix |
| --- | --- | --- | --- |
| TRM.all | 22,137 | Each tocochromanol phenotype | TRM using all 22,137 gene loci |
| TRM.cand | 111 | Each tocochromanol phenotype | TRM using 111 candidate gene loci |
| GRM+TRM.all | 22,137 | Each tocochromanol phenotype | GRM + TRM using all 22,137 gene loci |
| GRM+TRM.cand | 111 | Each tocochromanol phenotype | GRM + TRM using 111 candidate gene loci |
| Multi-trait GBLUP | 1-3 (phenotype dependent) | Each tocochromanol phenotype and transcripts of 1-3 gene loci | GRM |

<sup>a</sup> TRM, transcriptomic relationship matrix; GRM, genomic relationship matrix

Given that likely causal genes underlying large-effect JL-QTL controlling natural variation for the nine tocochromanol phenotypes have been identified and most of them appeared to affect trait variation through *cis*-acting effects on mRNA expression (Wu et al., 2022), we tested a multi-trait GBLUP approach in the 545 lines of the Ames panel that incorporated the expression-BLUE values of seven candidate causal genes (*dxs2*, *hppd1*, *hggt1*, *por1*, *por2*, *vte3*, and *vte4*) that underpinned JL-QTL having a percent phenotypic variance explained (PVE) that was ≥ 5% in the maize NAM panel (Supplemental Table S4). For each tocochromanol phenotype, both Box-Cox transformed tocochromanol-BLUEs and expression-BLUEs of the 1–3 large-effect candidate causal genes (PVE ≥ 5%) identified for the phenotype (Supplemental Table S4) were modeled as multiple response variables in the multi-trait GBLUP model (Table 1). The multi-trait GBLUP model, which included the GRM constructed from the 341,034 SNPs, was implemented in the MTM package in R (https://github.com/QuantGen/MTM/) with the same MCMC parameters used in the BGLR package for the GP with CV. As the transcript abundances and tocochromanol levels were measured on grain samples collected from different field experiments, residual covariance was assumed to be independent (‘DIAG’ option for the residual covariance). The predictive ability of each model was estimated with the identical CV approach used for GRM, TRM, and GRM+TRM models.

## RESULTS

### MK-GBLUP based on NAM JL-QTL

We leveraged multiple sources of prior biological information related to the accumulation of tocochromanols in maize grain for inclusion in GP models. Through the implementation of a MK-GBLUP modeling approach, 154 JL-QTL associated with the concentrations of nine tocochromanol grain phenotypes (from 12 to 21 JL-QTL per phenotype; Supplemental Table S2) in the maize NAM panel (Diepenbrock et al., 2017) were assessed for improving GP abilities of the same nine phenotypes within and between the maize Goodman and Ames panels. We constructed QTL-specific relationship matrices with SNPs sampled at three different genomic window sizes: ± 250 kb of JL peak SNP, ± 1 Mb of JL peak SNP, and JL-QTL SI. On average, the ± 250 kb, ± 1 Mb, and SI windows included 74, 304, and 2,602 SNPs per QTL for the nine phenotypes, respectively. As the width of SIs largely varied among QTL, window size based on SI ranged from 0.07 to 123.38 Mb (Supplemental Table S2). The generation of these matrices allowed us to also evaluate whether window size affects predictive abilities of MK-GBLUP models, given that both the precise causal variants underlying the JL-QTL and optimal size of windows are unknown.

The predictive abilities of the three MK-GBLUP models with different window sizes (± 250 kb, ± 1 Mb, and SI) were compared to each other using two whole-genome prediction models (GBLUP and BayesB) that have contrasting assumptions of genetic architecture (Figure 1 and Supplemental Table S5). Overall, moderate predictive abilities (range: *r* = 0.09 to 0.72; average: *r* = 0.42) were observed (Figure 1A) from the testing of five models on nine tocochromanol grain phenotypes in four prediction scenarios (from Ames to Goodman, from Goodman to Ames, within Ames, and within Goodman). Higher predictive abilities were, with one exception (αT, GBLUP), always observed within the Ames panel compared to within the Goodman panel, which is likely attributed in part to the 6-fold larger size of the Ames panel (*n* = 1,462) relative to the Goodman panel (*n* = 242). However, predictive abilities from Ames to Goodman were lower than that from Goodman to Ames in 28 out of 45 combinations of phenotypes and models. Interestingly, the difference in predictive abilities between the two prediction scenarios across the five models was more dependent on the tocochromanol phenotype, as predictive abilities for all five models from Ames to Goodman were higher than those from Goodman to Ames for δT, αT3, and δT3, while lower for γT, γT3, ΣT, ΣT3, and ΣTT3.

**Figure 1.**
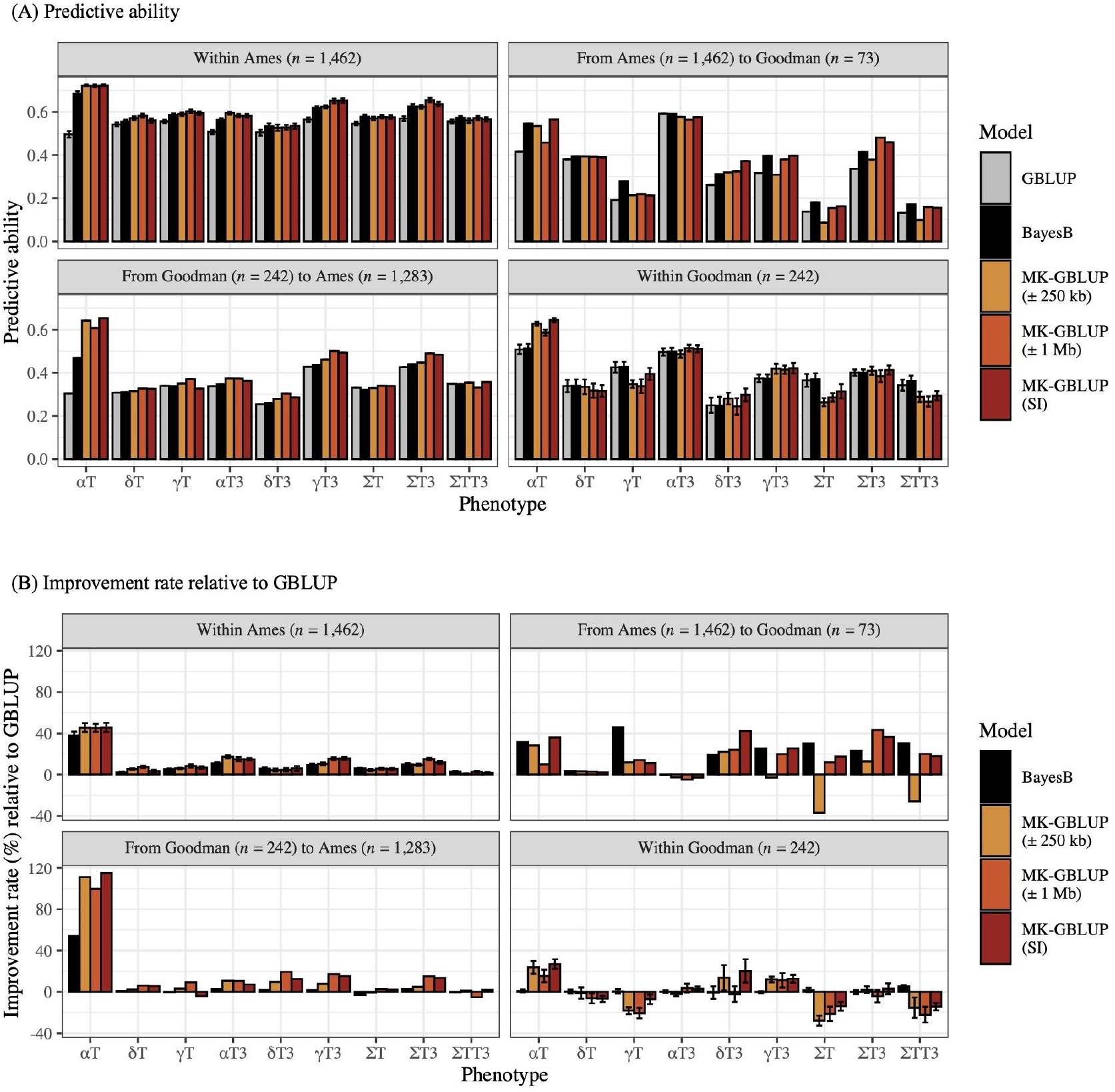
(A) Predictive abilities of tocochromanol grain phenotypes with genomic best linear unbiased prediction (GBLUP), BayesB, and multikernel GBLUP (MK-GBLUP) models within and between the Ames and Goodman panels, and (B) improvement rate (%) of BayesB and MK-GBLUP models relative to GBLUP. The MK-GBLUP model was tested with three genomic window sizes surrounding joint-linkage quantitative trait loci identified in the maize nested association mapping panel: ± 250 kb, ± 1 Mb, and support interval (SI). The nine predicted phenotypes are as follows: α-tocopherol (αT), δ-tocopherol (δT), γ-tocopherol (γT), α-tocotrienol (αT3), δ-tocotrienol (δT3), γ-tocotrienol (γT3), total tocopherols (ΣT), total tocotrienols (ΣT3), and total tocochromanols (ΣTT3). Predictive ability was expressed as the mean Pearson’s correlation (*r*) between predicted and observed values for each phenotype across 10 repetitions of five-fold cross-validation. The predictive ability of the GBLUP model was used as the baseline to calculate the improvement rate of the BayesB and MK-GBLUP models. Error bars represent one standard deviation from the mean predictive ability. As no subsampling occurred in either of the between panel prediction schemes, standard deviation was not calculated and therefore no error bars are displayed.

Given that GBLUP, which assumes polygenic genetic control, is arguably the most widely used model in GP, it was used as a baseline model in our study. When only considering the GBLUP results, the predictive abilities had an average of *r* = 0.39 across all prediction scenarios and phenotypes, ranging from *r* = 0.13 (for ΣTT3, prediction from Ames to Goodman) to *r* = 0.59 (for αT3, prediction from Ames to Goodman). Collectively, 78.5% of the evaluated MK-GBLUP or BayesB models had higher predictive abilities than the GBLUP models. With predictive abilities averaged across the nine phenotypes and four prediction scenarios, MK-GBLUP with SI window showed the best predictive ability (on average, improvement rate was +13.6%), followed by MK-GBLUP with ± 1 Mb window (+10.8%), BayesB (+10.2%), and MK-GBLUP with ± 250 kb window (+7.0%) relative to GBLUP. Notably, for αT, which is predominantly under the genetic control of the large-effect *vte4* gene (Diepenbrock et al., 2017), MK-GBLUP with SI widow showed the large improvement rates in all prediction scenarios (from +26.9% to +114.9%; Figure 1B) when compared to GBLUP.

The best performing prediction model of the five tested was dependent on the four prediction scenarios and nine tocochromanol phenotypes. Of the 36 combinations between the four prediction scenarios and nine tocochromanol phenotype, MK-GBLUP with SI window showed the best predictive ability for 12 combinations, MK-GBLUP with ± 1 Mb window for 11 combinations, BayesB for 10 combinations, and MK-GBLUP model with ± 250 kb window for two combinations. When these results were further examined, the MK-GBLUP models showed higher predictive abilities than BayesB for the tocochromanol phenotypes strongly controlled by *vte4* (αT and αT3) or *hggt1* (δT3, γT3, and ΣT3) (Diepenbrock et al., 2017). However, BayesB showed higher predictive abilities than the MK-GBLUP models for the two sum traits, ΣT and ΣTT3. The absolute value of the change in predictive ability was larger for the MK-GBLUP models than for the BayesB models when compared to GBLUP. Specifically, MK-GBLUP models showed -37.0% to +114.9% improvement rates, while BayesB had -2.9% to +53.9% improvement rates.

We evaluated whether the PVEs of 154 JL-QTL associated with tocochromanol grain phenotypes in the NAM panel are relatable to the predictive abilities observed between the Ames and the Goodman panels. To test this, the correlation between the PVEs for JL-QTL estimated in the NAM panel (Diepenbrock et al., 2017) and the predictive ability of each QTL-specific relationship matrix based on the ± 250 kb window (evaluated by using partial correlation) were calculated for both between-panel prediction scenarios. The PVEs and predictive abilities showed a strong positive Pearson’s correlation (*r* = 0.66 for the prediction from Ames to Goodman; *r* = 0.81 for the prediction from Goodman to Ames) when the correlation was calculated on the QTL-phenotype pairs with PVE ≥ 5%, regardless of the prediction scenarios. In contrast, a weaker positive correlation was found between them when PVE was < 5% (*r* = 0.02 for the prediction from Ames to Goodman; *r* = 0.35 for the prediction from Goodman to Ames). In further support of these findings, similar correlation values were obtained for both the MK-GBLUP models with ± 1 Mb or SI windows (Supplemental Table S6).

### Transcriptome-based prediction

Because the transcript abundances of genes may provide orthogonal regulatory information that cannot be provided by genetic marker data alone, we hypothesized that jointly modeling SNP marker genotypes and gene transcript abundances from developing grain would improve the predictive ability of the nine tocochromanol grain phenotypes. This hypothesis was tested on 545 lines of the Ames panel through the evaluation of five transcriptome-based prediction models that used different sets of transcripts with or without a GRM. Though not better than, the predictive ability based on all transcripts for the 22,137 gene loci (TRM.all model) was comparable to the GBLUP (GRM only) model, as the average improvement rate over the nine phenotypes was -3.8% (Figure 2 and Supplemental Table S7). When the TRM was built with only the 111 *a priori* candidate gene loci involved in the accumulation of tocochromanols in maize grain (TRM.cand model), the predictive ability decreased for all tocochromanol phenotypes except for αT, for which the predictive ability improved by 20.5% relative to the TRM.all model. In contrast, the average accuracy improvements obtained by the GRM+TRM.all and GRM+TRM.cand models were +1.1% and +7.6%, respectively, relative to the GBLUP model.

**Figure 2.**
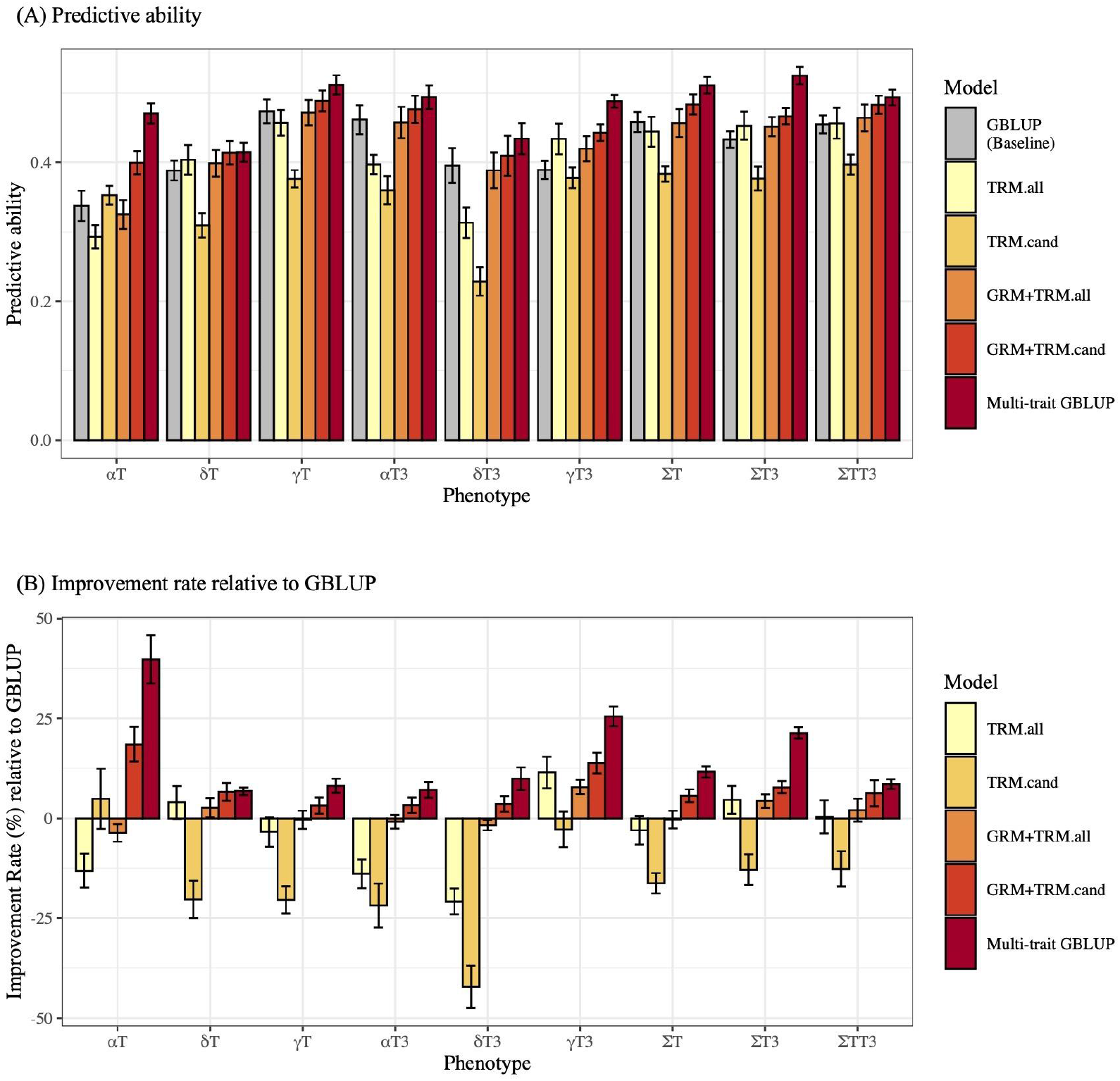
(A) Predictive abilities of tocochromanol grain phenotypes with the genomic best linear unbiased prediction (GBLUP) and transcriptome-based prediction models (Table 1) within the Ames panel, and (B) improvement rate (%) of transcriptome-based prediction models relative to GBLUP. The multi-trait GBLUP model includes a genomic relationship matrix (GRM) and uses the transcript abundance of one to three candidate causal genes underlying large-effect QTL with each tocochromanol phenotype as multiple response variables. The other four prediction models are as follows: a transcriptomic relationship matrix (TRM) incorporating all available genes (TRM.all), a TRM incorporating only 111 *a priori* candidate gene loci (TRM.cand), a GRM included with TRM.all (GRM+TRM.all), and a GRM included with TRM.cand (GRM+TRM.cand). The nine predicted phenotypes are as follows: α-tocopherol (αT), δ-tocopherol (δT), γ-tocopherol (γT), α-tocotrienol (αT3), δ-tocotrienol (δT3), γ-tocotrienol (γT3), total tocopherols (ΣT), total tocotrienols (ΣT3), and total tocochromanols (ΣTT3). Predictive ability was expressed as the mean Pearson’s correlation (*r*) between predicted and observed values for each phenotype across 10 repetitions of five-fold cross-validation. The predictive ability of the GBLUP model was used as the baseline to calculate the improvement rate of transcriptome-based prediction models. Error bars represent one standard deviation from the mean predictive ability.

We further investigated the genetics underlying improvements to predictive abilities of the two GRM+TRM models compared to the GBLUP model, because of our interest in how jointly modeling transcript abundances and SNP markers impacted model performance. By taking the ranking of absolute values of the regression coefficients for transcripts (*i.e.*, estimated effect sizes of transcripts) from the GRM+TRM.all model for each phenotype, we generated an importance score (Azodi et al., 2020) for each of the 22,137 gene loci. This allowed us to examine the relationship between the importance scores of 10 candidate causal genes identified to underlie QTL for controlling tocochromanol grain levels by Diepenbrock et al. (2017) that were included in the GRM+TRM.all model and their percent PVE estimated in the maize NAM panel. There were 21 gene-phenotype pairs with PVE ≥ 5%, and 18 of them had an importance score in the top 1% (Figure 3). The three exceptions were *vte3* for δT, *hppd1* for δT3, and *hggt1* for ΣTT3. Of those three, *vte3* and *hppd1* were not shown to be ceeQTL for any of the nine tocochromanol grain phenotypes in the maize NAM panel (Diepenbrock et al., 2017); therefore, it is reasonable that their importance score in the GRM+TRM.all model was not ranked in the top 1%.

**Figure 3.**
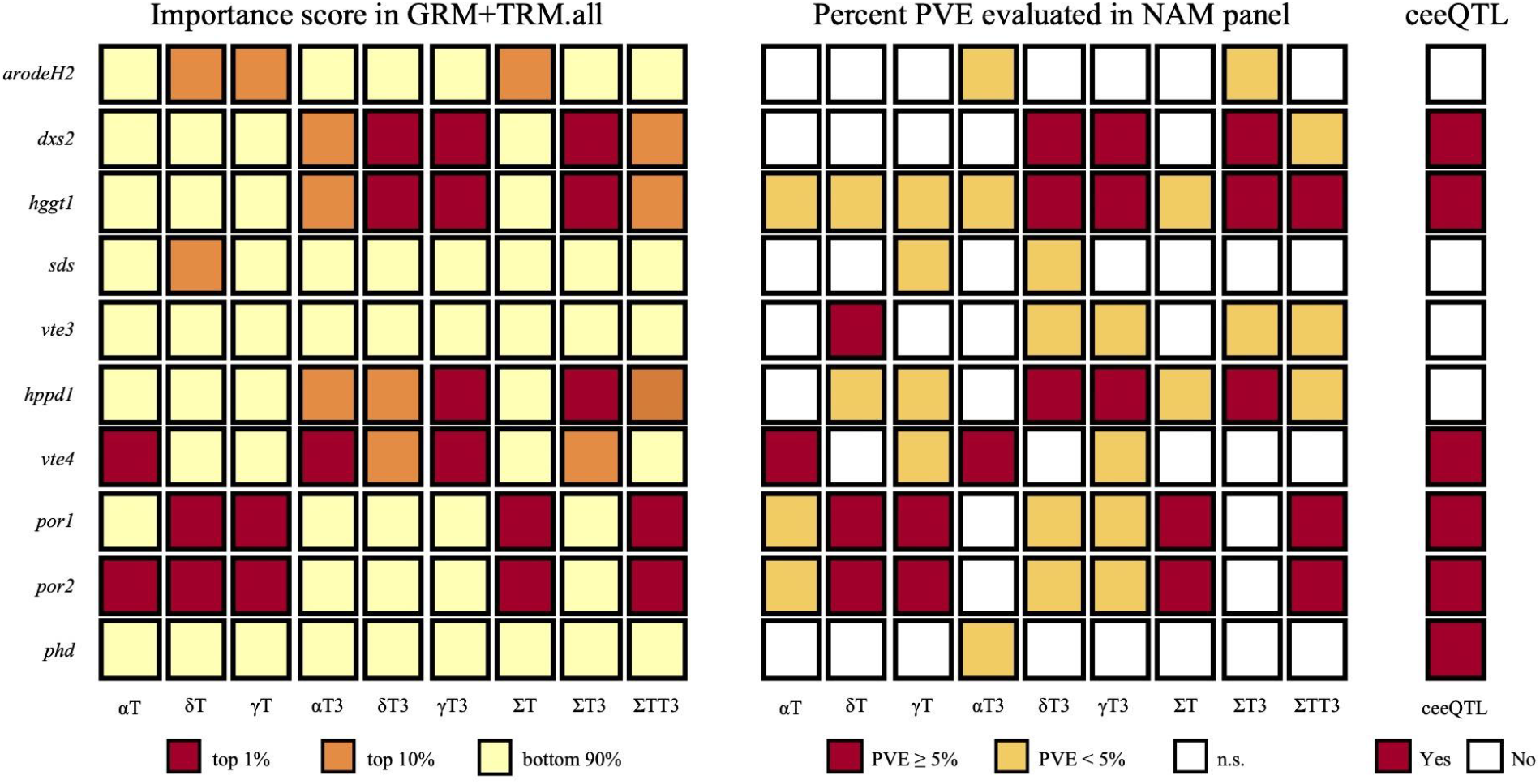
Importance scores of candidate causal genes identified for tocochromanol grain phenotypes in the U.S. maize nested association mapping (NAM) panel. The nine investigated phenotypes are as follows: α-tocopherol (αT), δ-tocopherol (δT), γ-tocopherol (γT), α-tocotrienol (αT3), δ-tocotrienol (δT3), γ-tocotrienol (γT3), total tocopherols (ΣT), total tocotrienols (ΣT3), and total tocochromanols (ΣTT3). For each phenotype, the importance scores of gene loci were determined by the ranking of absolute values of regression coefficients from the model that included a genomic relationship matrix (GRM) and transcriptomic relationship matrix (TRM) incorporating all available genes (GRM+TRM.all). Three (*snare*, *ltp*, and *fbn*) of the 13 candidate causal genes identified to associate with tocochromanols in the maize NAM panel by Diepenbrock et al (2017) were not included in the GRM+TRM.all model, because they did not pass quality control. In the left panel, a box for a gene with a top 1%, top 10% (but not in top 1%), or bottom 90% importance score is filled with a red, orange, or yellow color, respectively. In the middle panel, a box for a gene found to underlie a joint-linkage quantitative trait loci (JL-QTL) with a phenotypic variance explained (PVE) of ≥ 5% or < 5% in the NAM panel (Diepenbrock et al., 2017) is filled with a red or dark-yellow color, respectively. The box is filled with a white color if the gene (JL-QTL) was not significant (n.s.) for the phenotype in the NAM panel. In the right panel, a box for a gene designated as a correlated expression and effect QTL (ceeQTL) in the NAM panel (Diepenbrock et al., 2017) is filled with a red color.

We also performed a PCA of the regression coefficients for the 111 *a priori* candidate gene loci from the GRM+TRM.cand model for the six tocochromanol compounds and visualized the result in a two-dimensional PCA biplot by highlighting the 13 candidate causal genes identified for grain tocochromanols in the NAM and/or Ames panels (Diepenbrock et al., 2017; Wu et al., 2022) with at least a top 10% importance score for one or more of the six compounds (Figure 4). Most of the 13 genes were distant from the center in the biplot, indicating that these genes had large effects on a corresponding set of phenotypes as indicated by the directions of arrows for the six tocochromanol compounds. Indicative of its major role in the qualitative profile of tocopherols and tocotrienols, the *vte4* gene, which methylates γ/δ isoforms to produce α/β isoforms (Shintani & DellaPenna, 1998), had the largest positive effects on αT and αT3 among the 111 *a priori* candidate gene loci included in the prediction model. Supportive of earlier findings by Diepenbrock et al. (2017) and Wu et al. (2022), large positive effects of *por1* and *por2* were found for tocopherols (δT and γT), whereas tocotrienols (δT3 and γT3) were predominantly impacted by the large positive effects of *hggt1* and *dxs2*. When searching for other genes with the largest PC scores, we found that the gene with the most negative PC2 score, Zm00001d048736, codes for a protein with similarity to S-alkyl-thiohydroximate lyase (SUR1) in *Arabidopsis thaliana* (Supplemental Table S3). However, this gene had at best a top 10% importance score for δT, γT and ΣT among the 22,137 gene loci.

**Figure 4.**
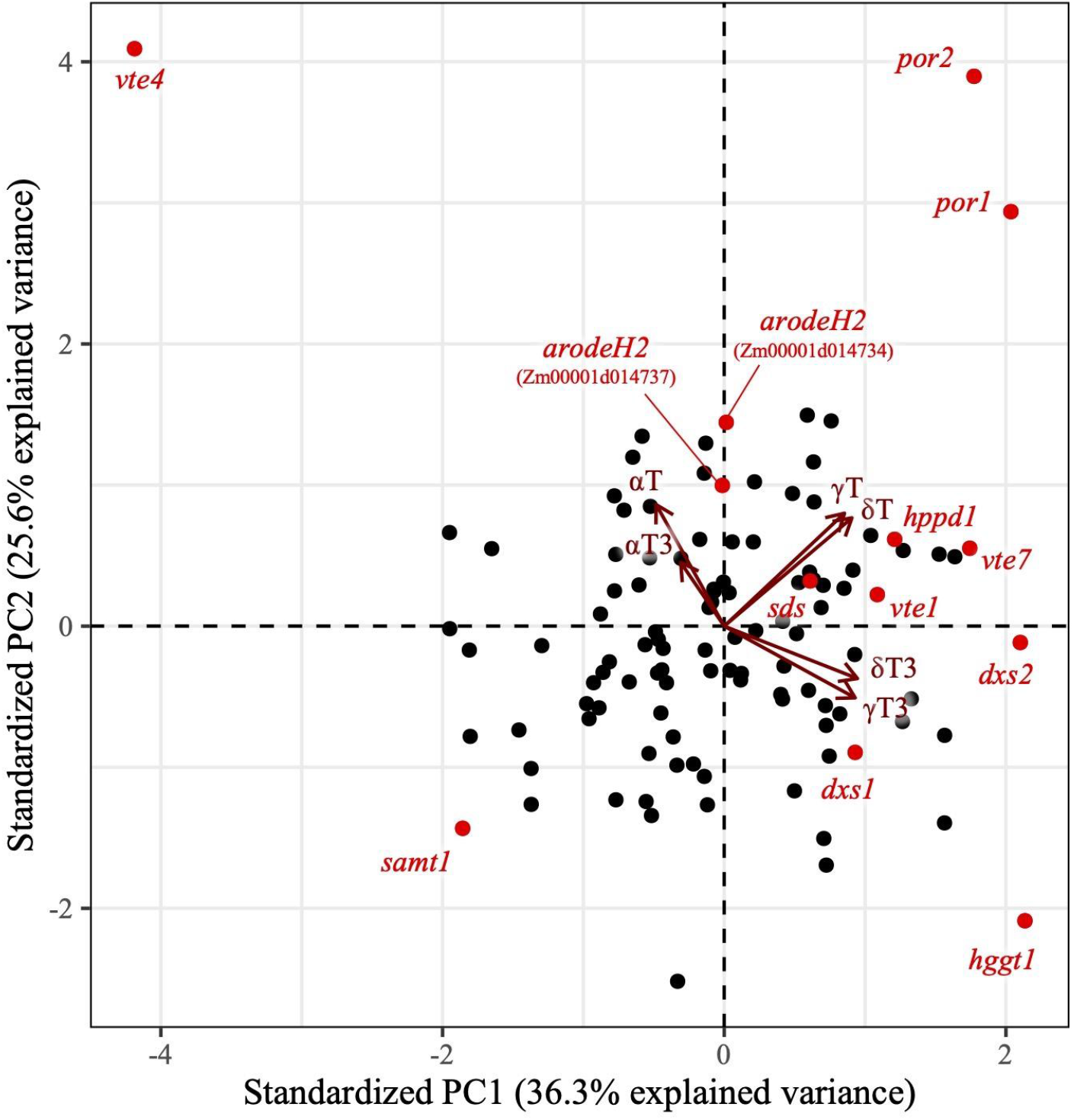
Biplot of both principal component scores (dots) and loadings (vectors) from a principal component analysis on the regression coefficients of 111 *a priori* candidate gene loci from the prediction of tocochromanols. The regression coefficients were generated from the single trait prediction of six tocochromanol compounds: α-tocopherol (αT), α-tocotrienol (αT3), γ-tocopherol (γT), γ-tocotrienol (γT3), δ-tocopherol (δT), and δ-tocotrienol (δT3). The model used for prediction included a genomic relationship matrix (GRM) and transcriptomic relationship matrix (TRM) incorporating only 111 *a priori* candidate gene loci (GRM+TRM.cand). Each dot represents one of the 111 *a priori* candidate gene loci, and each vector represents one of the six tocochromanol compounds. The candidate causal genes identified by Diepenbrock et al. (2017) and Wu et al. (2022) that were included in the GRM+TRM.cand model and had at least a top 10% importance score for one or more of the six compounds are labeled in red font.

The ranking of importance scores and PCA of regression coefficients results implicated the greater role of expression profiles for a few large-effect candidate causal genes to predict tocochromanol grain levels. To further explore these insights, we tested the predictive ability of a multi-trait GBLUP approach (Table 1). Depending on the tocochromanol phenotype, transcript abundances for 1–3 of seven large-effect candidate causal genes (PVE ≥ 5% in the NAM panel; *dxs2*, *hppd1*, *hggt1*, *por1*, *por2*, *vte3*, and *vte4*) were modeled with each tocochromanol phenotype as multiple traits (Supplemental Table S4). Of these genes, all of which but *hppd1* were ceeQTL (Figure 3). We found that, on average, predictive abilities from the multi-trait GBLUP model were +15.4% higher than those of the GBLUP model, ranging from +6.8% for δT to +39.8% for αT (Figure 2 and Supplemental Table S7). This average improvement rate of +15.4% over all nine phenotypes was higher than the average improvement rate of +13.3% based on the MK-GBLUP model with ± 1 Mb window (best model in this prediction scenario) compared to GBLUP when evaluated within the Ames panel. Although they are not directly comparable as the number of Ames accessions was different (1,462 lines in the MK-GBLUP analysis, whereas 545 lines in this transcriptome-based prediction analysis), our results suggest that transcript abundances of large-effect loci, especially if gene expression mediates the causal relationship, should be prioritized for improving the predictive ability of tocochromanol phenotypes.

## DISCUSSION

Genomic selection is a promising breeding approach particularly for phenotypes controlled by many small-effect loci (Lorenz et al., 2011; Heslot et al., 2015). From this perspective, tocochromanol grain phenotypes are an interesting breeding target in maize because these traits are predominantly controlled by large-effect loci, with a modest contribution from smaller effect loci (Diepenbrock et al., 2017). Therefore, a statistical model that can incorporate biological information related to previously identified large-effect QTL/genes is hypothesized to improve predictive abilities, as has been shown by Bernardo (2014) in simulation experiments and empirically by Rutkoski et al. (2014) for stem rust resistance in wheat. In this study, we leveraged existing biological data by including QTL-specific relationship matrices or transcript abundances in prediction models to evaluate whether they improve the predictive ability of tocochromanol levels in maize grain. Overall, we showed that these two data types increased predictive abilities and explored which genetic factors may have affected model performance.

In contrast to the genomic selection + *de novo* GWAS modeling approach of Spindel et al. (2016) that uses the training population to select significant GWAS markers for inclusion in trait prediction models, we implemented a multikernel modeling approach by constructing a local relationship matrix for QTL identified in a panel different from those used for trait prediction. When incorporating in GP models the 12–21 QTL-specific relationship matrices for NAM JL-QTL identified for tocochromanol grain phenotypes (Diepenbrock et al., 2017), variation in improvement rates were observed across phenotypes and prediction scenarios compared to the GBLUP model. Overall, we found that the improvement rate based on MK-GBLUP was higher when a larger amount of the phenotypic variance of the predicted phenotype was explained by a few large-effect genes. Furthermore, PVEs of JL-QTL in the NAM panel showed a strong positive correlation with between-population predictive abilities when JL-QTL had PVEs ≥ 5% (Supplementary Table S6). When the predictive ability of the MK-GBLUP model with SI interval was evaluated within the Ames population, the highest improvement was observed for αT (+45.8%), while the improvement rate was lowest for ΣTT3 (+1.9%) (Supplementary Table S5). The lower improvement rate for ΣTT3 is likely to be explained by the complex genetic architecture of ΣTT3, which is a summation of six different tocochromanol compounds. Indeed, the largest PVE observed in the NAM panel for a gene shown to genetically control ΣTT3 was 12.3% for *por2*, whereas *vte4* explained 48.2% of the phenotypic variance for αT (Diepenbrock et al., 2017). Altogether, we showed that modeling large-effect QTL with a MK-GBLUP modeling approach is valuable for improving the predictive ability of tocochromanol levels in maize grain.

Although the MK-GBLUP model improved predictive abilities for the majority of tocochromanol phenotypes in most prediction scenarios, low or even negative improvement may be observed as the effect of a QTL/gene can be population- and/or environment-specific (de Roos et al., 2009; Windhausen et al., 2012; Widener et al., 2021). In our study, the MK-GBLUP models showed lower predictive abilities than the GBLUP model irrespective of the window size when predicting δT and γT within the Goodman panel. Even though the *por1* and *por2* genes have PVEs ≥ 5% (range: 5.9–17.5%) for δT and γT in the maize NAM panel, these loci were not significantly associated with these or any other tocochromanol grain phenotype via GWAS in the Goodman panel (Kremling et al., 2019). However, *por2*—the larger effect of the two *por* loci—was detected by GWAS for grain δT and γT levels with ∼1,500 lines of the Ames panel, but the GWAS only detected an association of *por1* with ΣT (Wu et al., 2022). Therefore, large-effect causal variants at both *por* loci in the Goodman panel may be at a markedly lower frequency relative to the Ames panel, which could result in the observed negative improvements when predicting δT and γT in the Goodman panel (Heslot et al., 2012; Scutari et al., 2016). These results illustrate that the transferability of QTL from biparental populations to diversity panels for use in prediction models is not always exact even for large-effect QTL. To improve the low predictive ability for these two phenotypes in the Goodman panel, it could be worthwhile to apply training set optimization approaches that were recently tested by Tibbs-Cortes et al. (2022) for prediction of grain tocochromanols in exotic-derived maize.

When large-effect QTL or genes are identified by linkage analysis or GWAS, the most popular method is to include the peak SNP markers as fixed covariates in the GBLUP model (Bernardo, 2014; Spindel et al., 2016). In our study, however, we implemented a multikernel approach by constructing a local relationship matrix for each JL-QTL. This was based on the assumption that the peak SNP at each JL-QTL is not necessarily in strong LD with causal variants across different mapping panels, thus it is better to use multiple SNPs within a QTL interval to adequately circumscribe causal variants across multiple panels. Therefore, we tested three window sizes (± 250 kb, ± 1 Mb, and SI), but found that predictive abilities were largely robust to the choice of window size. Although the amount and pattern of LD decay depends on the population and genomic region, LD decays very rapidly (average *r^2^* of 0.2 within ∼1–10 kb) in diverse groups of maize germplasm (Romay et al., 2013). Therefore, it is likely that the narrowest window (± 250 kb) tested was sufficient to encompass causal variants underlying JL-QTL. Whether the optimization of window sizes for the LD patterns of each JL-QTL would lead to further improvement is an open question, but such continued investigation is limited by not knowing the true causal variants.

We tested five different models that included transcriptome data to predict tocochromanol grain phenotypes. Consistent with the findings of Azodi et al. (2020) and Hershberger et al. (2022), prediction models that incorporated both a GRM and TRM were generally superior to models that only used one of them. More interestingly, when our tested models included a GRM, it was better to include transcript abundances for a subset of candidate genes prioritized for their purported role in the biosynthesis of tocochromanols in maize grain rather than transcriptome-wide abundances from all 22,137 gene loci (GRM+TRM.all model). Of the two *a priori* candidate gene-targeted prediction models tested, the highest predictive ability was achieved with a multi-trait GBLUP model that had a tocochromanol phenotype and transcript abundances of only 1–3 genes determined to underlie large-effect JL-QTL (PVE ≥ 5%) in the NAM panel as multiple response variables. The second of the two models, which had a GRM and TRM constructed from the transcript abundances of 111 *a priori* candidate gene loci (GRM+TRM.cand model), also outperformed the GRM+TRM.all model. Collectively, to improve predictive ability based on transcripts and SNPs, prioritization of genes according to *a priori* knowledge (*e.g.*, biosynthetic pathway or known causal genes) on the target phenotype is suggested when possible, but the gain in predictive ability is likely to be dependent on the trait and population.

When prioritizing candidate or causal genes for inclusion in transcriptome-based prediction models, it could be better to preferentially consider genes that influence phenotypic variation through a mechanism mediated by *cis*-expression QTL (*cis*-eQTL). All five genes (*por1*, *por2*, *hggt1*, *dxs2*, and *vte4*) underlying large-effect (PVE ≥ 5%) JL-QTL and designated as ceeQTL in the NAM panel (Diepenbrock et al., 2017) showed a top 1% importance score in transcriptome-based prediction for multiple gene-phenotype pairs. What biological mechanisms could be responsible for such high importance scores for these five genes? In the Ames panel, Wu et al. (2022) detected *dxs2*, *por1*, *por2*, *hggt1*, and *vte4* by TWAS, thus demonstrating a strong relationship between grain tocochromanol and gene expression variation. With the exception of *dxs2* which was not detected by GWAS, Wu et al. (2022) implicated *cis*-acting variants as causal for co-localized GWAS and *cis*-eQTL association signals at the *por1*, *por2*, *hggt1*, and *vte4* loci. Given the interrelatedness of these findings, there exists an opportunity to extend computational approaches such as Camoco (Schaefer et al., 2018) for integrating co-expression networks with GWAS, TWAS, and eQTL mapping results to better prioritize candidate causal genes that further increase the predictive ability of transcriptome-based prediction models for nutritional grain phenotypes.

## CONCLUSIONS

To accelerate the genetic improvement of grain tocochromanol levels in maize biofortification breeding programs, we posited that harnessing biologically-relevant information would boost the predictive abilities of GP models for these nutritionally important phenotypes. By implementing a MK-GBLUP modeling approach that leveraged JL-QTL associated with tocochromanol grain phenotypes in the maize NAM panel, we improved predictive abilities for most of the nine tocochromanol phenotypes in all four prediction scenarios evaluated within and between the maize Ames and Goodman panels through capturing both large- and small-effect loci. Our results suggest that the inclusion of QTL-specific relationship matrices for large-effect loci in the GBLUP model is optimal for phenotypes that have an oligogenic architecture that includes several major QTL such as tocochromanols (Diepenbrock et al., 2017) and carotenoids (Diepenbrock et al., 2021) in maize grain, given that the GBLUP model is limited by its inability to have a few markers with large effects (Meuwissen et al., 2001; de Los Campos et al., 2013; Gianola, 2013).

Through testing of transcriptome-based prediction models within the Ames panel, we showed that the largest gains in predictive abilities were achieved with a multi-trait GBLUP model that fitted a tocochromanol phenotype and the transcript abundances of 1–3 genes determined to underlie large-effect JL-QTL (PVE ≥ 5%) as multiple response variables. Most of these large-effect genes were implicated as ceeQTL with *cis*-acting variants responsible for variation in grain tocochromanols (Diepenbrock et al., 2017; Wu et al., 2022). This implies a higher importance of purportedly causal genes with large expression-based effects when predicting tocochromanols in maize grain. Considering the decreasing cost of RNA sequencing and availability of a maize practical haplotype graph (Valdes Franco et al., 2020), the predictive ability of our transcriptome-based prediction approaches can be likely improved upon by including Bayesian networks (dos Santos et al., 2020) to model the expression profiles of several candidate causal genes collected from multiple kernel developmental time points and enable the between-population transfer of imputed expression effects associated with *cis* haplotypes (Giri et al., 2021).

## ACKNOWLEDGMENTS

We thank Matt Baseggio, Elise Albert, and other current and past members of the Gore, DellaPenna, Buell, and Yu labs for their efforts in pollination, harvest, and sample preparation. This research was supported by the National Science Foundation (IOS-1546657 to J.Y., C.R.B, D.D.P., and M.A.G.); the National Institute of Food and Agriculture; the USDA Hatch under accession numbers 1013641 and 1023660 (M.A.G.), and 1021013 (J.Y.); Cornell University startup funds (M.A.G.); and HarvestPlus (M.A.G.). L.T.C. was supported by the National Science Foundation Graduate Research Fellowship Program (NSF GRFP grant no. 1744592). This study was also made possible by the support of the American People provided to the Feed the Future Innovation Lab for Crop Improvement through the United States Agency for International Development (USAID) (M.A.G.). The contents are the sole responsibility of the authors and do not necessarily reflect the views of USAID or the United States Government. Program activities are funded by the United States Agency for International Development (USAID) under Cooperative Agreement No. 7200AA-19LE-00005.

## AUTHOR CONTRIBUTIONS

Ryokei Tanaka: Conceptualization; Data curation; Formal analysis; Methodology; Visualization; Writing – original draft; Writing-review & editing.

Di Wu: Data curation; Formal analysis; Methodology; Writing – review & editing.

Xiaowei Li: Data curation; Formal analysis; Methodology; Writing – review & editing.

Joshua C. Wood: Data curation; Formal analysis; Methodology; Writing – review & editing.

Laura E. Tibbs-Cortes: Data curation; Formal analysis; Methodology; Investigation; Writing – review & editing.

Maria Magallanes-Lundback: Data curation; Investigation; Writing – review & editing.

Nolan Bornowski: Data curation; Formal analysis; Writing – review & editing.

John P. Hamilton: Data curation; Formal analysis; Writing – review & editing.

Brieanne Vaillancourt: Data curation; Formal analysis; Writing – review & editing.

Xianran Li: Formal analysis; Writing – review & editing.

Nicholas T. Deason: Data curation; Investigation; Writing – review & editing.

Gregory R. Schoenbaum: Data curation; Investigation; Writing – review & editing.

Jianming Yu: Funding acquisition; Resources; Supervision; Writing – review & editing.

C. Robin Buell: Funding acquisition; Resources; Supervision; Writing – review & editing.

Dean DellaPenna: Funding acquisition; Resources; Supervision; Writing – review & editing.

Michael A. Gore: Conceptualization; Funding acquisition; Methodology; Project administration; Resources; Supervision; Writing-review & editing

## CONFLICT OF INTEREST

The authors declare no conflict of interest.

## SUPPLEMENTAL MATERIAL

**Supplemental Table S1.** Summary of the 1,704 maize inbred lines used in this study

**Supplemental Table S2.** Summary of the evaluated multikernel GBLUP (MK-GBLUP) models

**Supplemental Table S3.** Genomic information of the 126 *a priori* candidate gene loci used for transcriptome-based prediction

**Supplemental Table S4.** Large-effect genes [percent phenotypic variance explained (PVE) ≥ 5%] that were associated with tocochromanol grain phenotypes in the U.S. maize nested association mapping (NAM) panel (Diepenbrock et al., 2017). Depending on the phenotype, transcript abundances of 1-3 large-effect genes were used as response variables with a tocochromanol phenotype in the multi-trait GBLUP model.

**Supplemental Table S5.** Predictive abilities and improvement rates (%) based on GBLUP, BayesB, and multikernel GBLUP (MK-GBLUP) models

**Supplemental Table S6.** Pearson’s correlation between phenotypic variance explained (PVE) of joint-linkage quantitative trait loci (JL-QTL) identified for tocochromanol grain phenotypes in the U.S. maize nested association mapping (NAM) panel and predictive ability of the multikernel GBLUP (MK-GBLUP) models

**Supplemental Table S7.** Predictive abilities and improvement rates (%) based on GBLUP and transcriptome-based prediction models

## DATA AVAILABILITY STATEMENT

All raw 3′ mRNA-seq data are available from the NCBI Sequence Read Archive under BioProject PRJNA643165. Supplementary SNP genotype, tocochromanol grain phenotype, and developing grain transcriptome data sets are available on CyVerse (https://de.cyverse.org/data/ds/iplant/home/shared/GoreLab/dataFromPubs/Tanaka_TocochromanolPrediction_2022). All code is available on Github (https://github.com/GoreLab/Tocochromanol_Prediction).

## Abbreviations

BLUE: best linear unbiased estimator
BLUP: best linear unbiased predictor
ceeQTL: correlated expression and effect QTL
CV: cross-validation
eQTL: expression QTL
GBLUP: genomic best linear unbiased prediction
GBS: genotyping-by-sequencing
GP: genomic prediction
GRM: genomic relationship matrix
GWAS: genome-wide association studies
HPLC: high-performance liquid chromatography
IBS: identity-by-state
JL: joint linkage
LD: linkage disequilibrium
MCMC: Markov chain Monte Carlo
MK-GBLUP: multikernel genomic best linear unbiased prediction
NAM: nested association mapping
PVE: phenotypic variance explained
PCA: principal component analysis
QTL: quantitative trait loci
SNP: single-nucleotide polymorphism
TRM: transcriptomic relationship matrix
TWAS: transcriptome-wide association studies
SI: support interval
αT: α-tocopherol
αT3: α-tocotrienol
γT: γ-tocopherol
γT3: γ-tocotrienol
δT: δ-tocopherol
δT3: δ-tocotrienol
ΣT: total tocopherols
ΣT3: total tocotrienols
ΣTT3: total tocochromanols

## Notes

### Competing Interest Statement

The authors have declared no competing interest.

